# Calcium-dependent transcriptional profiles of human pancreatic islet cells reveal functional diversity in islet subpopulations

**DOI:** 10.1101/2023.09.08.556709

**Authors:** Ji Soo Yoon, Shugo Sasaki, Jane Velghe, Katarina Zosel, Michelle Y. Y. Lee, Helena Winata, Cuilan Nian, Francis C. Lynn

## Abstract

**Aims/hypothesis:** Pancreatic islets depend on cytosolic calcium to trigger the secretion of glucoregulatory hormones and regulate the transcription of genes important for the response to stimuli. To date, there has not been an attempt to profile calcium-regulated gene expression in all islet cell types. Our aim was to construct a large single-cell transcriptomic dataset from human islets exposed to conditions that would acutely induce or inhibit intracellular calcium signalling, while preserving biological heterogeneity.

**Methods:** We exposed intact human islets from three donors to the following conditions: (1) 2.8 mM glucose; (2) 25 mM glucose and 40 mM KCl to maximally stimulate calcium signalling; and (3) 25 mM glucose, 40 mM KCl and 5 mM EGTA (calcium chelator) to inhibit calcium signalling, for 1 hour. We sequenced 43,909 cells from all islet cell types, and further subsetted the cells to form an endocrine cell-specific dataset of 32,486 cells expressing *INS*, *GCG*, *SST* or *PPY*. We compared transcriptomes across conditions to determine the differentially expressed calcium-regulated genes in each endocrine cell type, and in each endocrine cell subcluster of alpha and beta cells.

**Results:** Based on the number of calcium-regulated genes, we found that each alpha and beta cell cluster had a different magnitude of calcium response. We also showed that a polyhormonal cluster expressing *INS, GCG*, and *SST* is defined by calcium-regulated genes specific to this cluster. Finally, we identified the gene *PCDH7* from the beta cell clusters that had the highest number of calcium-regulated genes, and showed that cells expressing cell surface PCDH7 protein have enhanced glucose-stimulated insulin secretory function.

**Conclusions:** Here we use our single-cell dataset to show that human islets have cell-type-specific calcium-regulated gene expression profiles, some of them specific to subpopulations. In our dataset, we identify *PCDH7* as a novel marker of beta cells having an increased number of calcium-regulated genes and enhanced insulin secretory function.

**Data availability:** A searchable and user-friendly format of the data in this study, specifically designed for rapid mining of single-cell RNA sequencing data, is available at https://lynnlab.shinyapps.io/Hislet_2023/. The raw data files are available at NCBI Gene Expression Omnibus (GSE196715).

## Introduction

The pancreatic islets of Langerhans is a glucoregulatory micro-organ composed of insulin (INS)-secreting beta (β) cells, glucagon (GCG)-secreting alpha (α) cells and somatostatin (SST)-secreting delta (δ) cells. Like most electrically excitable secretory cells, islets require calcium influx to secrete glucoregulatory hormones. However, intracellular calcium leads to other processes in islets, including transcription^1^. As recently reviewed^1^, two main pathways regulate calcium-dependent transcription: (1) the calmodulin-dependent protein kinase (CAMK)/cAMP response element-binding protein (CREB) pathway^1–6^; and (2) the calcineurin (CaN)/nuclear factor of activated T cells (NFAT) pathway^1,7^. Due to the complexities of calcium signalling and the ready availability of mouse and human β cell lines, most studies have focused on calcium signalling specifically in β cells^5,8,9^.

Using single-cell RNA sequencing (scRNA-seq), it is now possible to study multiple cell types in islets simultaneously. A number of human islet scRNA-seq datasets have focused on how diabetes alters the islet transcriptome, identifying rare cell types, and coupling function to transcriptomes^10–15^. Here, we used scRNA-seq to identify rapidly responding calcium-regulated genes in islet cell types. We generated an adult human islet dataset from three donors, using islets exposed to three experimental conditions. Our scRNA-seq dataset is available as a user-friendly web tool for studying islet heterogeneity and transcriptional response to stimuli.

## Methods

### Human islets

Human islets were isolated by the University of Alberta Islet Research or Clinical Cores as described (dx.doi.org/10.17504/protocols.io.x3mfqk6. Accessed 20-01-2021). Details of donor metrics and functional data are available at https://www.epicore.ualberta.ca/IsletCore/ and are summarised in Table 1.

**Table 1.**
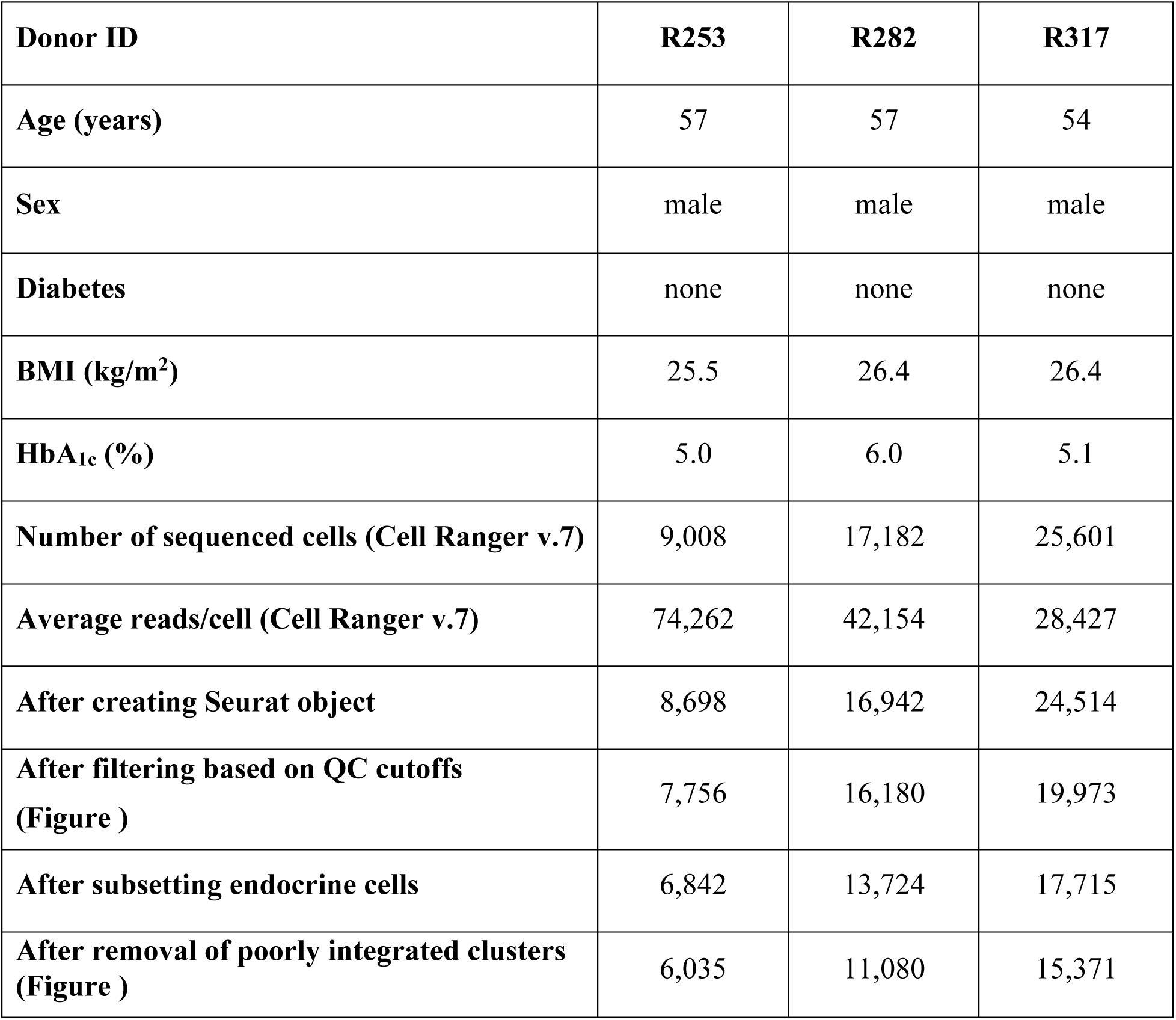
Human islet donor information and cell numbers during key analysis steps.

| **Donor ID** | **R253** | **R282** | **R317** |
| --- | --- | --- | --- |
| **Age (years)** | 57 | 57 | 54 |
| **Sex** | male | male | male |
| **Diabetes** | none | none | none |
| **BMI (kg/m<sup>2</sup>)** | 25.5 | 26.4 | 26.4 |
| **HbA<sub>1c</sub> (%)** | 5.0 | 6.0 | 5.1 |
| **Number of sequenced cells (Cell Ranger v.7)** | 9,008 | 17,182 | 25,601 |
| **Average reads/cell (Cell Ranger v.7)** | 74,262 | 42,154 | 28,427 |
| **After creating Seurat object** | 8,698 | 16,942 | 24,514 |
| **After filtering based on QC cutoffs (Figure )** | 7,756 | 16,180 | 19,973 |
| **After subsetting endocrine cells** | 6,842 | 13,724 | 17,715 |
| **After removal of poorly integrated clusters (Figure )** | 6,035 | 11,080 | 15,371 |

### Human islet stimulation and dispersion

Islets were handpicked into CMRL 1066 medium (VWR, USA; CA45001-114) and incubated overnight at 37°C in a humidified CO2 incubator. The next morning, 450 islets per donor were handpicked into three wells (150 islets/well) of a 12-well plate containing Krebs-Ringer Bicarbonate HEPES (KRBH) medium (2.8 mM glucose) and incubated at 37°C for 1 hour. Next, the islets were incubated for 1 hour in a new well containing KRBH under one three experimental conditions: Low (2.8 mM glucose); Positive (25 mM glucose, 40 mM KCl); or Negative (25 mM glucose, 40 mM KCl, 5 mM EGTA). Islets were washed with PBS (Mg^2+^/calcium-free) containing 0.5 mM EDTA, then dispersed for 12–15 minutes in 200 µL of 0.25% trypsin/EDTA at 37°C. Trypsinisation was quenched with 25% FBS (1 ml; in PBS). Cells were centrifuged for 3 minutes at 200 *g*, the supernatant fraction was removed, and cells were resuspended in 300 µL of PBS with 2% FBS. Cells were filtered through a 40 µm cell strainer (Corning, USA; 352340). Cells were counted prior to centrifugation and resuspension in PBS at 1000 cells/µL for scRNA-seq.

### NanoString gene profiling assay

50 islets from each donor batch were used to assess islet quality by profiling expression of 132 human islet genes (Supplementary Table 1). Gene expression was measured using nCounter prep kits and nCounter SPRINT profiler according to manufacturer’s instructions (NanoString, USA).

### scRNA-seq

libraries were generated with 10x Genomics (USA) Chromium single-cell 3’ reagent kits according to manufacturer’s instructions. Version 2 reagent kits were used for donors R253 and R282, and Version 3 was used for donor R317. Each experimental condition was labelled as a sample, with a total of three samples per donor. Each donor library pool was sequenced twice using the Illumina NextSeq500 with NextSeq 500/550 High Output v2 or v2.5 75-cycle kits (Illumina, USA; FC-404-2005). Libraries were sequenced using the following cycle parameters: 28 (Read 1), 50 (Read 2), 8 (i7), 0 (i5) for donors R253 and R282 libraries. Sequencing for donor R317 used 56 cycles (Read 2) as recommended by 10x Genomics (USA).

### Analysis of scRNA-seq data

Sequenced data files were initially processed using Cell Ranger V.7.0.1 (10X Genomics) software. FASTQ files were generated from sequencing data using *cellranger mkfastq*, then aligned to reference human genome GRCh38 to generate gene counts using *cellranger count*. Following Cell Ranger pipelines, additional quality control and filtering of cells was performed using publicly available R package Seurat V.4.0.4^16^. Seurat was used to filter out genes that were not expressed in at least 3 cells, cells that expressed more than 20% mitochondrial genes, and cells that expressed more than 6000 genes. Seurat was also used for integration of the 3 donor datasets, clustering, visualizing, and finding differentially expressed genes (DEG), including calcium-regulated genes.

For identifying calcium-regulated genes, all cells were identified by their corresponding donor ID and experimental condition (Low, Positive, or Negative). Cell types and cluster identities were added to the metadata and new Seurat Objects were produced for each cell type. The dataset was parsed by cell type and three subsets were generated to contain two experimental conditions in each subset. This process was repeated in order to generate two-way comparisons for all combinations of conditions in each cell type. Final calcium-regulated genes in each cluster per cell type were identified as significantly expressed genes in either the Positive or Negative condition with an adjusted p-value less than 0.05.

RNA Velocity analysis was conducted to infer the direction of transcriptional dynamics using the scVelo Python package v0.2.1^17^ (Theis lab, Germany; https://github.com/theislab/scvelo). Count matrices of unspliced and spliced abundances were obtained from fastq files using the loompy/kallisto pipeline (loompy v3.0.6, kallisto v0.46.0, https://linnarssonlab.org/loompy/kallisto/index.html). L files across patient and condition were merged, and scVelo was ran following the pipeline for the endocrine pancreas dataset (https://scvelo.readthedocs.io/about/). Metadata and Uniform Manifold Approximation and Projection (UMAP) embeddings were exported from our processed Seurat object (for only endocrine cells) and merged with the splicing information. Genes with less than 20 spliced and unspliced counts were filtered out, and counts were normalized by the total initial counts in each cell. The first- and second-order moments for each cell were computed using its 30 nearest neighbours on the kNN graph in PC space (30 PCs). Velocity vectors were estimated by solving a stochastic model of transcriptional dynamics and the expected mean direction of all possible cell transitions on the kNN graph were calculated. Cosine correlations between velocity vectors and possible cell transitions were obtained in high dimensional space to generate probabilities for all potential cell transitions. The resulting matrix was used to project the velocities into lower-dimensional UMAP space for visualization.

### Immunofluorescence staining

Briefly, 100–200 human islets per donor were fixed in 4% paraformaldehyde for 1 hour, embedded in 2% agarose, paraffin-embedded, and sectioned at 5 µm. The sections were de-paraffinized, rehydrated, blocked and incubated with primary antibodies at 4°C overnight in PBS containing 5% horse serum. Sections were washed and incubated with secondary antibodies for 1 hour at 22°C. Sections were imaged using a Leica SP8 confocal microscope. A list of antibodies used for immunofluorescence staining can be found in the Supplementary Methods.

### RNAscope fluorescence *in situ* hybridisation

5 μm sections of embedded human islets or human pancreas biopsies were probed for human *INS* and *GCG* mRNA using RNAscope fluorescent multiplex v2 kit according to manufacturer’s instructions (ACDbio, USA). Probes for human *INS* (313571-C2) and *GCG* (556741-C3) were incubated with human islet sections. Opal dyes 520 and 650 (Perkin-Elmer, USA; FP1487001KT, FP1496001KT) at 1:100 dilution were used to develop *INS* and *GCG* signals, respectively, and DAPI was used as the nuclear stain. Sections were imaged using the SP8 confocal microscope (Leica, Germany).

### FACS and reaggregation of protocadherin 7-positive cells

Human islets were dispersed as above, washed with PBS, and incubated with rabbit AlexFluor647-labelled anti-protocadherin 7 (PCDH7) antibodies (Bioss Antibodies, USA; bs-11085R-A647) on ice for 30 minutes. Cells were washed with PBS, filtered (40 µm), resuspended in 500 µL of 2% FBS in PBS, and sorted using a BD FACSAria IIu (BD Biosciences, USA). Sorted PCDH7^+^ and PCDH7^−^ cells were plated onto a Corning Elplasia 96-well round bottom ultra-low attachment microcavity microplate (Corning, USA; 4442) at 80 aggregates/well (1000 cells/aggregate) and cultured in CMRL medium for 48 hours prior to determining glucose-stimulated insulin secretion (GSIS).

### GSIS assay

Islets from donors R366, R367, R369, H2330, H2337, and H2338 were incubated in KRBH with 2.8 mM glucose for 1 hour, then sequentially stimulated with 2.8 mM glucose, 16 mM glucose, and 2.8 mM glucose with 40 mM KCl in KRBH for 1 hour each at 37°C. Human C-peptide ELISA kits (Mercodia, Sweden; 10-1136-01) were used and stimulation index was calculated by normalising to values at 2.8 mM glucose.

### Quantitative PCR

PCDH7+ and - cells were sorted into TRIzol. RNA was isolated, DNase-treated with the Turbo DNA-free kit (Ambion, USA; AM1907), and reverse-transcribed with SuperScript^TM^ III (ThermoFisher, USA; 18080093). TaqMan quantitative PCR (qPCR) was performed to assess expression levels of *PCDH7*, *INS*, *ERO1B*, *NKX6-1* and *SLC30A8,* all normalized to reference gene *TBP.* A complete list of primers and probes used for qPCR can be found in the Supplementary Methods.

### Statistical analyses

All DEG were determined as those with an adjusted *p* value (*p*adjusted) less than 0.05 using a non-parametric Wilcoxon rank sum test. Student’s *t* test was performed on qPCR data, with significance defined as *p*<0.05. All statistical analyses were performed using Rstudio or GraphPad Prism software v.8.0.1.

## Results

### ScRNA-seq identification of human islet cell types

We generated islet single-cell libraries from three healthy male, BMI- and age-matched donors (Table 1). Islet quality was assessed prior to library generation by comparing islet donor gene expression (panel of 132 genes) to that in islets of 107 healthy people (Supplementary Table 1). To study islet cell calcium-regulated genes, we stimulated islets for 1 hour under the following conditions: (1) “Low” glucose to stimulate α cells and inhibit β cells; (2) “Positive”, a high glucose/high KCl stimulus to depolarise cells and stimulate calcium influx; and (3) “Negative”, a control condition with high glucose/high KCl stimulus and calcium chelator EGTA (Fig. 1a). The short stimulation time was to ensure the detection of only the most robust and acutely calcium-regulated genes.

**Figure 1.**
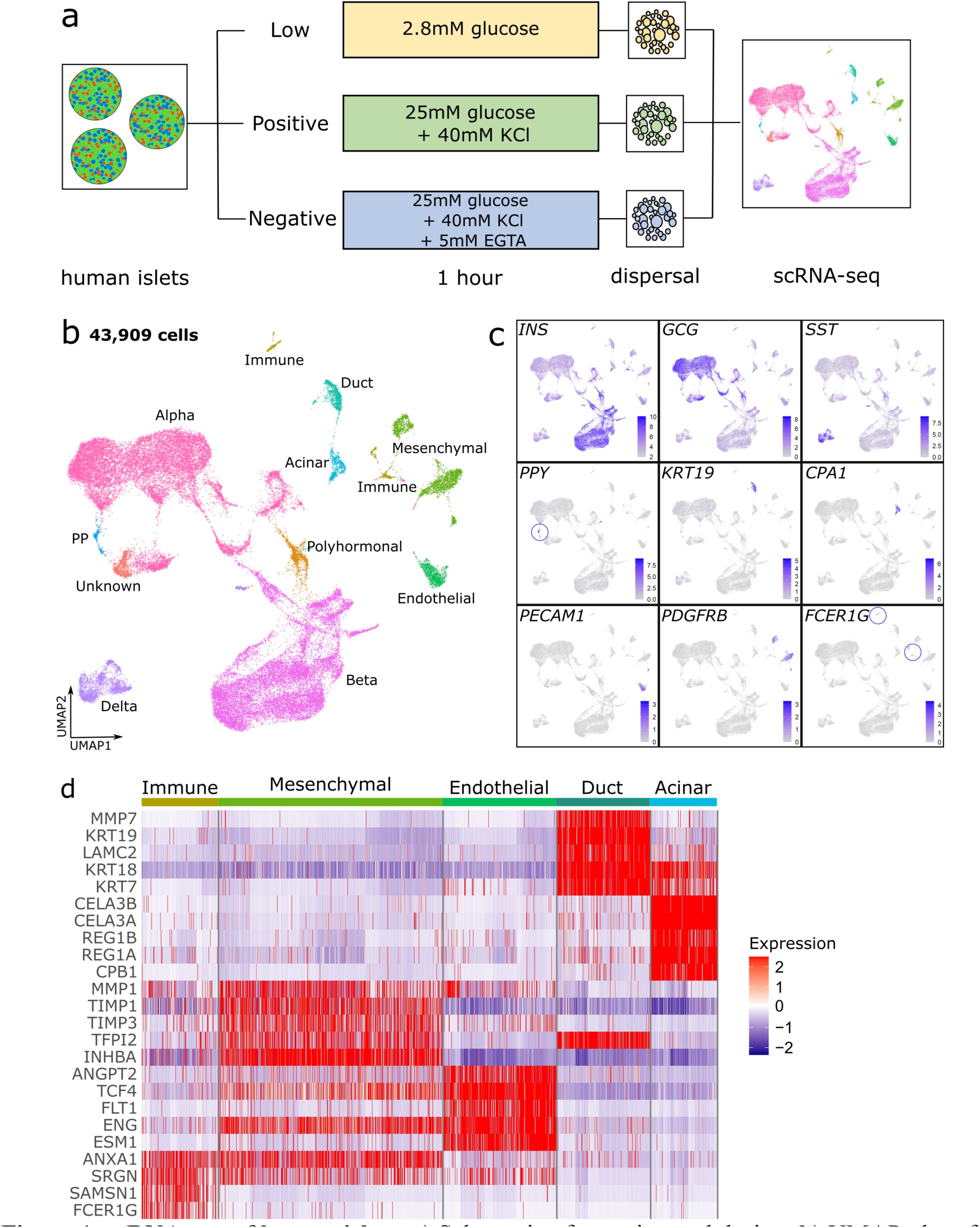
scRNA-seq of human islets. **a**) Schematic of experimental design. **b**) UMAP plot of 43,909 cells, clustered and labelled by cell type. **c**) Feature plots showing expression of known marker genes for each cell type. The immune cell clusters with high and specific *FCER1G* expression is highlighted in circles. **d**) Heatmap of top five DEG for each non-endocrine cell type vs all other cell types, with expression shown as log fold change.

Of the 51,791 sequenced cells, 84.8% passed the quality control thresholds, resulting in 43,909 cells as 30 clusters (Supplementary Fig. 1a). The remaining clusters showed similar proportions of Low, Positive and Negative cells, implying that clustering was minimally influenced by experimental conditions and that existing biological heterogeneity was preserved (Supplementary Fig. 1b). This supported our goal of designing experimental conditions that would allow us to investigate calcium-regulated genes and islet heterogeneity with the same dataset.

In the 30 clusters, there were 37,436 endocrine and 6,473 non-endocrine cells (Supplementary Fig. 1c,d). We determined cluster identities using known marker genes for each cell type: *INS* for β cells, *GCG* for α cells, *SST* for δ cells, *PPY* for Pancreatic Polypeptide (PP) cells, *KRT19* for duct cells, *CPA1* for acinar cells, *PECAM1* for endothelial cells, *PDGFRB* for mesenchymal cells, and *FCER1G* for immune cells^18^ (Fig. 1b,c). In addition to known cell types, there was a “polyhormonal” cluster expressing *INS*, *GCG,* and *SST,* and an “unknown” cluster that did not express robust levels of any of the known marker genes (Fig. 1b,c). For further validation of non-endocrine cell types, we carried out an unbiased DEG analysis, and found that *INHBA* and *TIMP1* expression were restricted to mesenchymal cells, while *PLVAP* and *ESM1* were endothelial cell marker genes (Fig.1d and Supplementary Fig.1e). Finally, we detected ghrelin (*GHRL*)-expressing cells located near the PP and α cells, but these cells did not form a whole cluster by themselves, presumably due to their low cell number (Supplementary Fig. 1f).

### α and β cells have distinct cluster-specific gene expression profiles

We subsetted and reclustered 38,281 α, β, δ, PP, and polyhormonal cells for further analysis (Supplementary Table 2). After reclustering, we removed clusters that were poorly integrated across donors, leaving 32,486 cells in 15 clusters (Fig 2a). Cluster identification was performed by expression of the key endocrine cell type marker genes *INS, GCG, SST,* and *PPY* (Supplementary Fig. 2a) and was confirmed by additional DEG analysis across the cell types (Supplementary Table 3). While δ, PP, and polyhormonal cells remained as single clusters, there were five clusters within α cells and seven clusters within β cells, which were named α1–α5 and β1–β7, respectively (Fig. 2a, Supplementary Fig. 2b). Overall, the dataset was now composed of 43% α cells, 46% β cells, 6.5% δ cells, 3.8% polyhormonal cells, and less than 1% PP cells.

**Figure 2.**
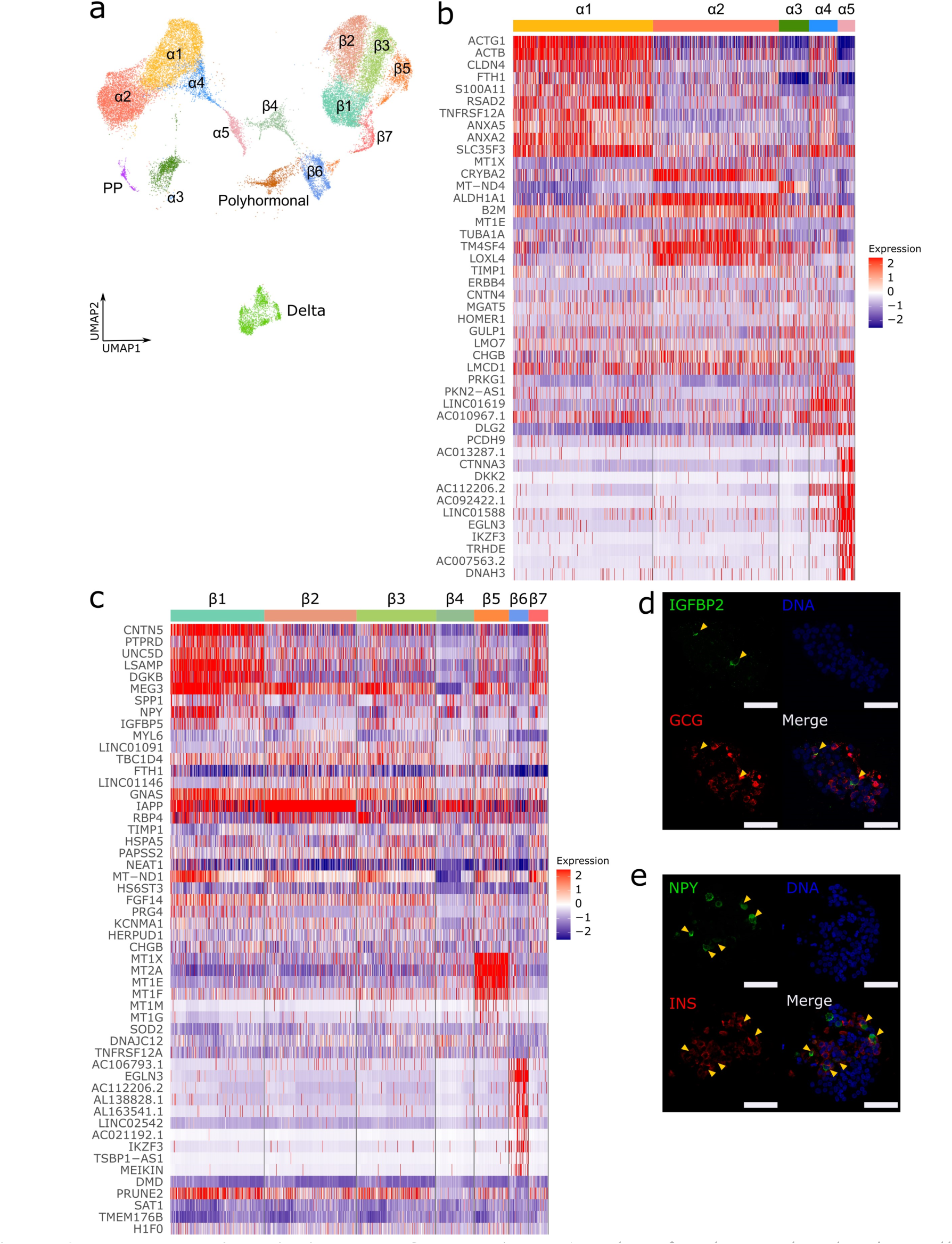
Heterogeneity within α and β cells. **a**) UMAP plot of reclustered endocrine cells. **b**, **c**) Heatmap of top ten DEG in α cell clusters (**b**) and β cell clusters (**c**), with expression shown as log fold change. **d**) Representative images of human islets immunostained for insulin like growth factor binding protein 2 (IGFBP2, green) and GCG (red). Arrowheads indicate IGFBP2^+^ GCG^+^ cells. **e**) Representative images of human islets immunostained for INS (red) and neuropeptide Y (NPY, green). Arrowheads indicate NPY^+^ INS^+^ cells. Scale bars, 50 μm.

To examine cluster heterogeneity among α and β cells and potentially uncover new cluster-specific marker genes, we performed DEG analysis across the clusters (Supplementary Tables 4,5). Within the α cell clusters, α1 and α4 expressed similar panels of marker genes, including *RSAD2* and *CLDN4* (Fig. 2b, Supplementary Fig. 2c). Cluster α2 was enriched for some α cell marker genes such as *ALDH1A1*, *CRYBA2*, *TM4SF4*^12^, and *LOXL4* (Fig. 2b, Supplementary Fig. 2c). Notably, none of the detected marker genes for cluster α3 seemed to be particularly robust. Despite its small cell numbers, the α5 cluster DEG list showed robust marker genes, including *CTNNA3* and *CHGB* (Fig. 2b, Supplementary Fig. 2c). Finally, the gene *IGFBP2* was expressed in a small proportion of all α cell clusters and showed high α cell specificity when compared with other cell types (Supplementary Fig. 2c). To confirm this, we immunostained human islet sections and found IGFBP2 protein was restricted to a small number of GCG-positive α cells (Fig. 2d).

Within β cells, the β1 cluster had elevated levels of genes such as *CNTN5, SPP1, NPY* and *IGFBP5* (Fig. 2c, Supplementary Fig. 2c). Immunostaining for NPY, which has been suggested to mark immature β cells^19^, showed expression in a subset of INS-expressing β cells (Fig. 2e). Cluster β2 showed the highest expression levels of *IAPP* among all clusters (Fig. 2c, Supplementary Fig. 2c). High protein levels of human islet amyloid polypeptide are associated with a pathological islet, β cell maturity or β cell dysfunction^20,21^. Similar to α3, clusters β3 and β4 did not show robustly elevated levels of any DEG, but cluster β4 uniquely expressed reduced levels of *KCNMA1*, which encodes for an α-subunit of a calcium-sensitive potassium channel (Supplementary Fig. 2c). Cluster β5 was enriched for metallothionein genes such as *MT2A*, *MT1X* and *MT1E*, suggesting increased protective capacity against oxidative stress^22^ (Fig. 2c, Supplementary Fig. 2c). Notably, clusters β4 and β6 were characterised by low expression of key β cell maturation markers such as *PDX1*, *UCN3* and *ERO1B* and could comprise transcriptionally immature β cells, such as virgin β cells^23^ (Supplementary Fig. 3a). In addition, β4 and β6 also showed low expression of several metabolically critical genes such as *GCK, ATP2A2,* and *G6PC2,* indicating a potentially different metabolic profile compared to other β cell clusters (Supplementary Fig. 3a).

To determine whether any α or β cell clusters were ‘metabolically immature’, we examined average expression of genes involved in the tricarboxylic acid (TCA) cycle, oxidative phosphorylation (OxPhos) and glycolysis in each cluster. In all gene panels, cluster α5 consistently showed overall lower expression, while α2 had elevated expression (Supplementary Fig. 3b). In β cells, cluster β4 showed reduced expression of TCA cycle and glycolysis genes but elevated expression of OxPhos genes, while β6 showed consistently reduced expression across all gene panels (Supplementary Fig. 3c). It is likely that the transcriptional immaturity suggested by lower expression of β cell identity genes in clusters β4 and β6 is linked to an immature metabolic state. These results imply a small proportion of metabolically unique α and β cells exist in the adult human islet.

### Identification of calcium-regulated gene sets in α cells

We next focused on identifying calcium- and glucose-regulated genes by paired comparisons of transcriptomes from different conditions. Within each cluster of a certain cell type, we identified calcium-regulated genes as those that had a higher expression in either the Positive or Negative condition when the two transcriptomes were compared. In the same manner, in the Low vs Positive comparison, genes that were significantly elevated in either condition would be glucose-induced. An ideal calcium-regulated gene would be expressed at lower levels in low glucose (Low), higher levels in response to depolarisation and calcium signalling (Positive), and at reduced levels when calcium signalling is inhibited by EGTA (Negative) (Fig. 3a).

**Figure 3.**
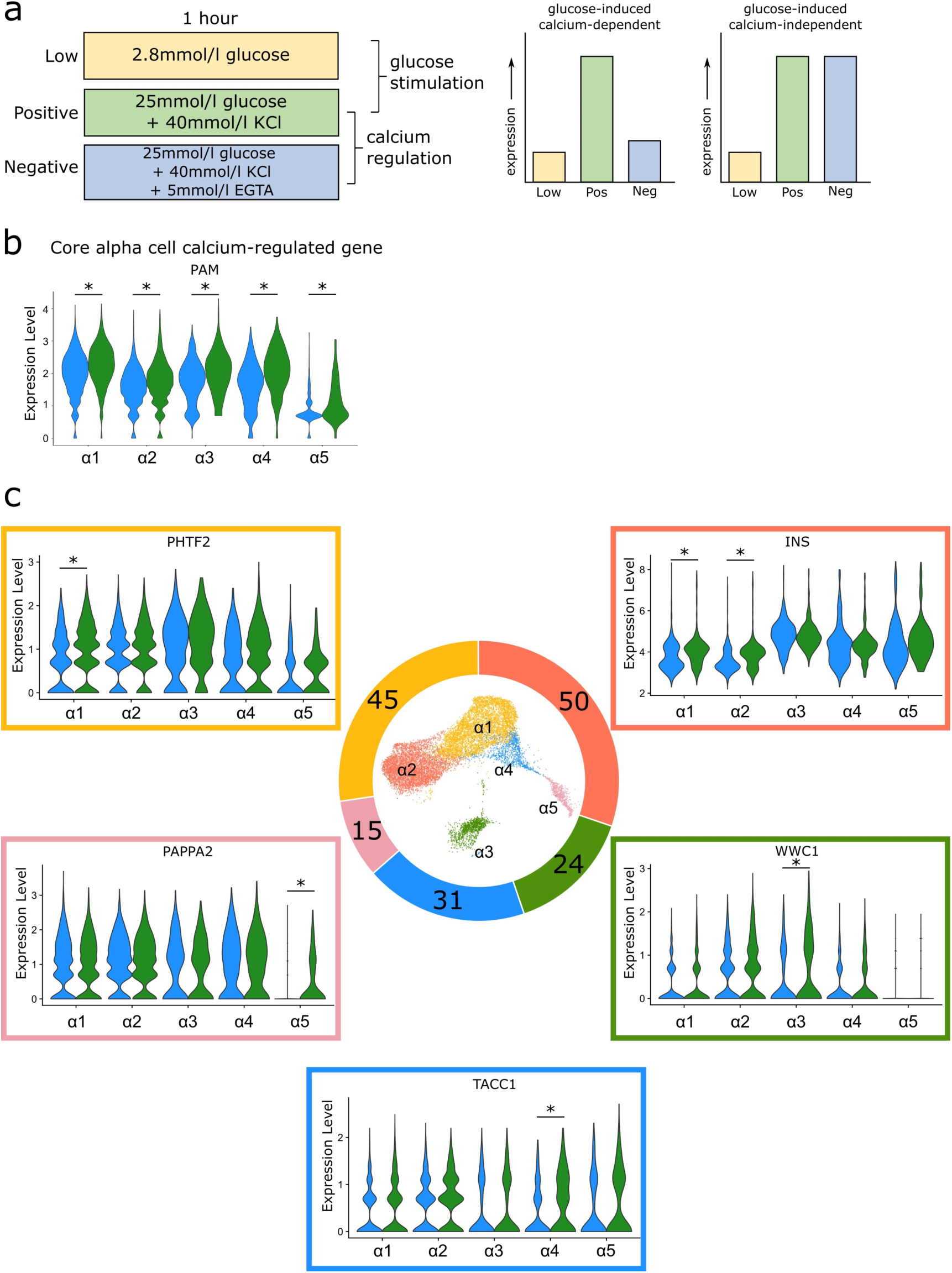
Calcium-regulated genes in α cells. **a**) Schematic for the strategy of identifying calcium-regulated and glucose-regulated genes, and example gene expression profiles across conditions for ideal calcium-dependent and calcium-independent genes. **b**) Split-violin plots showing the Core calcium-regulated gene *PAM* in α1–α5 clusters. **c**) Total number of calcium-regulated genes per cluster shown in the donut pie chart, with representative gene expression levels in Positive (green) and Negative (blue) conditions. \**p*adjusted<0.05.

In α cells, only the enzyme-encoding gene *PAM* was detected as calcium-regulated in all five α cell clusters (Fig. 3b). Cluster α2, which expressed the highest levels of α cell marker genes, had the highest number of cluster-specific calcium-regulated genes (Fig. 3c, Supplementary Table 6). α1 and α2 also had the highest number of calcium-regulated genes overall compared to all other α cell clusters (Fig. 3c, Supplementary Table 6). Interestingly, both α1 and α2 regulated typical β cell genes such as *IAPP* and *INS* (Fig. 3c, Supplementary Table 6). Overall, there were five calcium-regulated genes common to four out of five clusters, and nine calcium-regulated genes common to three out of five clusters (Supplementary Table 6). By the absolute number of calcium-regulated genes, we suspect that α1 and α2 are the most calcium-responsive clusters, while α5 may have a blunted calcium response. Since α cells secrete glucagon under low glucose conditions, we also compared Low and Negative conditions. In contrast to the comparison between Positive and Negative conditions, the Low vs Negative comparison showed only the gene *PTEN* as a commonly regulated gene between all five clusters (Supplementary Fig. 4a). As expected, most clusters had more genes expressed at higher levels in the Low condition, with the exception of α3 (Supplementary Fig. 4b). Once again, α5 showed a minimal response, with only one gene unique to the cluster (Supplementary Fig. 4b). From these results, we conclude that while most α cells show a transcriptional response to calcium and glucose, a small subset (α5) is less responsive or slower to respond within 1 hour of stimulation.

### Identification of calcium-regulated gene sets in β and δ cells

Unlike α cells, β and δ cells secrete hormones when ambient glucose levels are high, and have similar intracellular mechanisms downstream of glucose uptake^24^. Therefore, we focused on first identifying the calcium-regulated genes by comparing Positive vs Negative conditions. Cluster β6 had only three detectable calcium-regulated genes: *ELMO1, MT-ND3,* and *ZNF331* (Fig. 4a). These three genes were calcium-regulated in all seven β cell clusters and made up the “Core” list of β cell calcium-regulated genes. This meant that β6 did not have any cluster-specific calcium-regulated genes, further supporting the idea that β6 could be dysfunctional or immature compared with other β cell subtypes. The remaining six clusters had four common calcium-regulated genes in addition to the Core list: *C2CD4B, ELL2, SIK2,* and *SIK3* (Supplementary Fig. 5a). There were also several known immediate early genes (IEG) shared between four or five clusters, including *NR4A1* and *NR4A2*^25^, with β1-β3 having the highest degree of overlap (Supplementary Fig. 5b). β1-β3 also expressed the highest number of calcium-regulated genes (Fig. 4b, Supplementary Table 6), including known activity-regulated genes like *IAPP* and *NPAS4*^26,27^ (Fig. 4b, Supplementary Fig. 5b). In summary, β1-β3 are the most calcium-responsive β cell clusters, and β cells have an overall more homogeneous calcium-regulated profile than α cells, based on the number of Core calcium-regulated genes.

**Figure 4.**
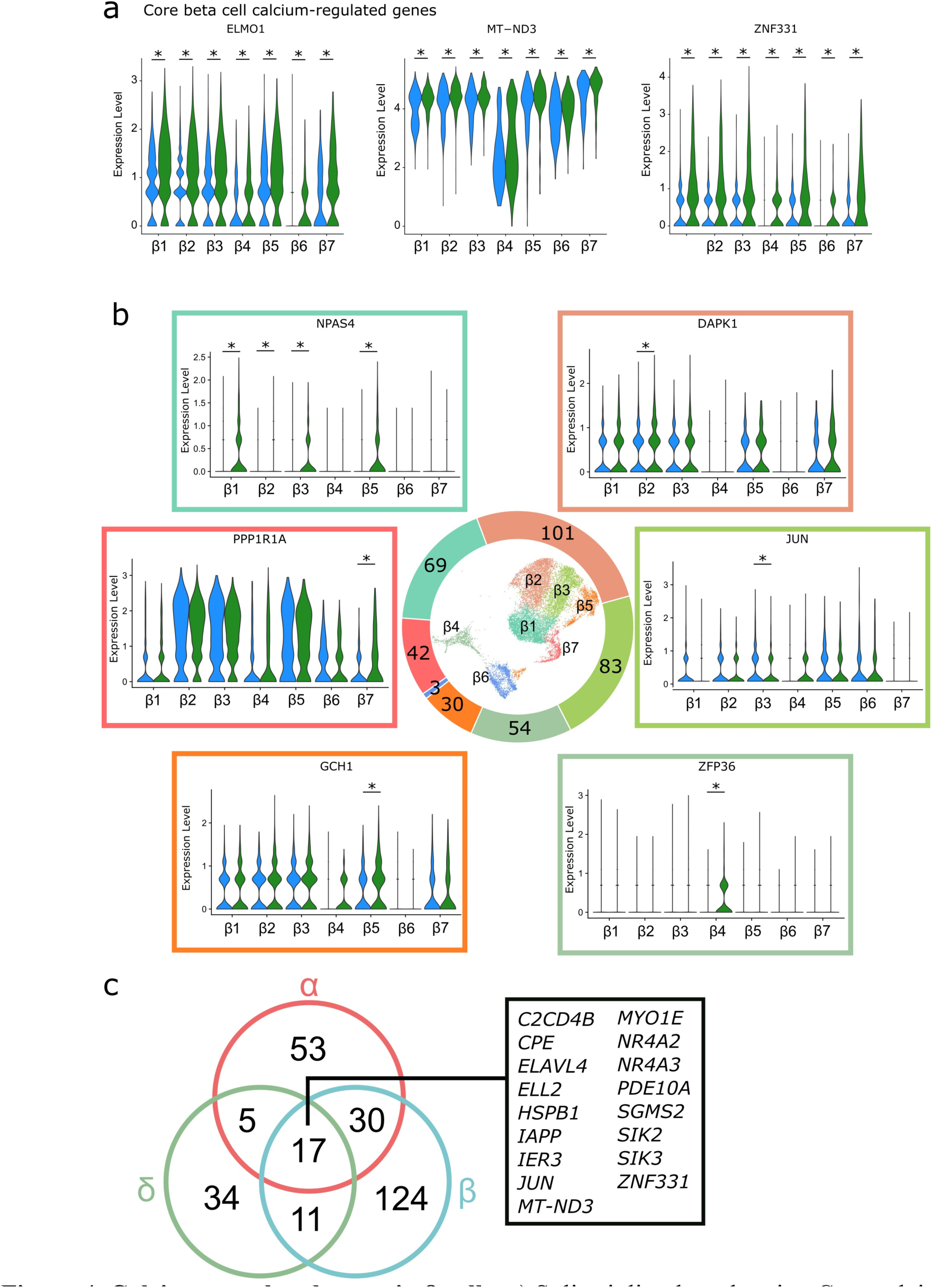
Calcium-regulated genes in β cells. **a**) Split-violin plots showing Core calcium-regulated genes in β1–β7 clusters. **b**) Total number of calcium-regulated genes per cluster shown in the donut pie chart, with representative gene expression levels in Positive (green) and Negative (blue) conditions. \**p*adjusted<0.05. **c**) Venn diagram of shared and unique calcium-regulated genes between α, β, and δ cells.

Next, we compared the Low and Positive conditions to determine the glucose-regulated profile for β cells. Once again, β6 seemed to be unresponsive, with no detected glucose-regulated genes (Supplementary Table 6). In contrast, most β cell clusters had more than 50 glucose-regulated genes, with the majority being glucose-induced (Supplementary Table 6). In at least two clusters, activity-regulated genes such as *NR4A1*, *NR4A2*, *IER3*, and *NPAS4* were all glucose-induced and calcium-regulated (Supplementary Fig. 5c, Supplementary Table 6). However, not all glucose-induced genes were calcium-regulated in the same clusters, and vice versa. For example, *FOS* was glucose-induced in β1-β5, but it was calcium-regulated in only three clusters (Supplementary Fig. 5d). Conversely, genes such as *C2CD4B* and *ELL2,* which were calcium-regulated in almost all clusters, were only glucose-induced in 1 or 2 clusters (Supplementary Fig. 5d). From these results, we conclude that not all glucose-regulated genes are calcium-regulated, and purely calcium-regulated genes are detectable using the method of pairwise cross-conditional comparisons.

Next, we carried out the same pairwise conditional comparisons in δ cells. Δ cells had fewer calcium-regulated genes than α or β cells, with 67 calcium-regulated genes and 95 glucose-regulated genes detected (Supplementary Table 6). Of these, 34 calcium-regulated genes were unique to δ cells, including *C4orf48, KLF10, NPHP4,* and *TMEM51* (Fig. 4c, Supplementary Fig. 6a). Likewise, 44 out of the 95 glucose-regulated genes were unique to δ cells (Supplementary Fig. 6b). From comparing all calcium-regulated genes in α, β, and δ cells, I found 17 genes that were calcium-regulated in all three cell types (Fig. 4c). One of these was *IAPP*, even though it is thought to be specific to adult β cells (Fig. 4c). Similarly, *INS* was calcium-regulated in δ cells and α cells, but not to any significant degree in β cells (Supplementary Table 6). Comparison of glucose-regulated genes across the three cell types also showed that *IAPP* was one of the 24 genes that are commonly glucose-regulated. Again, not all calcium-regulated genes are glucose-regulated, but genes such as *CPE, HSPB1, IER3, SIK2,* and *SIK3* were calcium-regulated and glucose-regulated in α, β, and δ cells (Supplementary Fig. 6b). The majority of genes were specific to an endocrine cell type, showing that adult human islet cell types have their own unique calcium-regulated transcriptional response.

### Polyhormonal cells in the adult human islet have a unique gene expression profile compared to α and β cells

One cluster in the dataset co-expressed *INS, GCG*, and to a lesser degree, *SST* as well (Fig. 5a). Comparing *INS, GCG*, and *SST* levels across all endocrine cell types shows that polyhormonal cells express lower levels of all three genes compared to α, β, and δ cells, but the proportion of cells expressing *INS, GCG*, and *SST* is almost 100% (Fig. 5a). This suggests that the polyhormonal cluster is not composed of distinct groups of *INS-*expressing cells and *GCG*-expressing cells, but true polyhormonal cells expressing multiple combinations of these hormone-encoding genes. We do not believe these are scRNA-seq artifacts composed of multiplets (multiple cells encased in a single droplet and barcoded as one cell), for the following reasons: First, comparing the number of genes and transcripts per cell across all endocrine cells in our dataset shows that polyhormonal cells do not express significantly higher (doubled or tripled) numbers of genes and/or transcripts, which would be most likely for a multiplet (Supplementary Fig. 7a,b). Second, levels of housekeeping genes such as *ACTB* and *RPLP0* are comparable between polyhormonal cells and other endocrine cells (Supplementary Fig. 7c).

**Figure 5.**
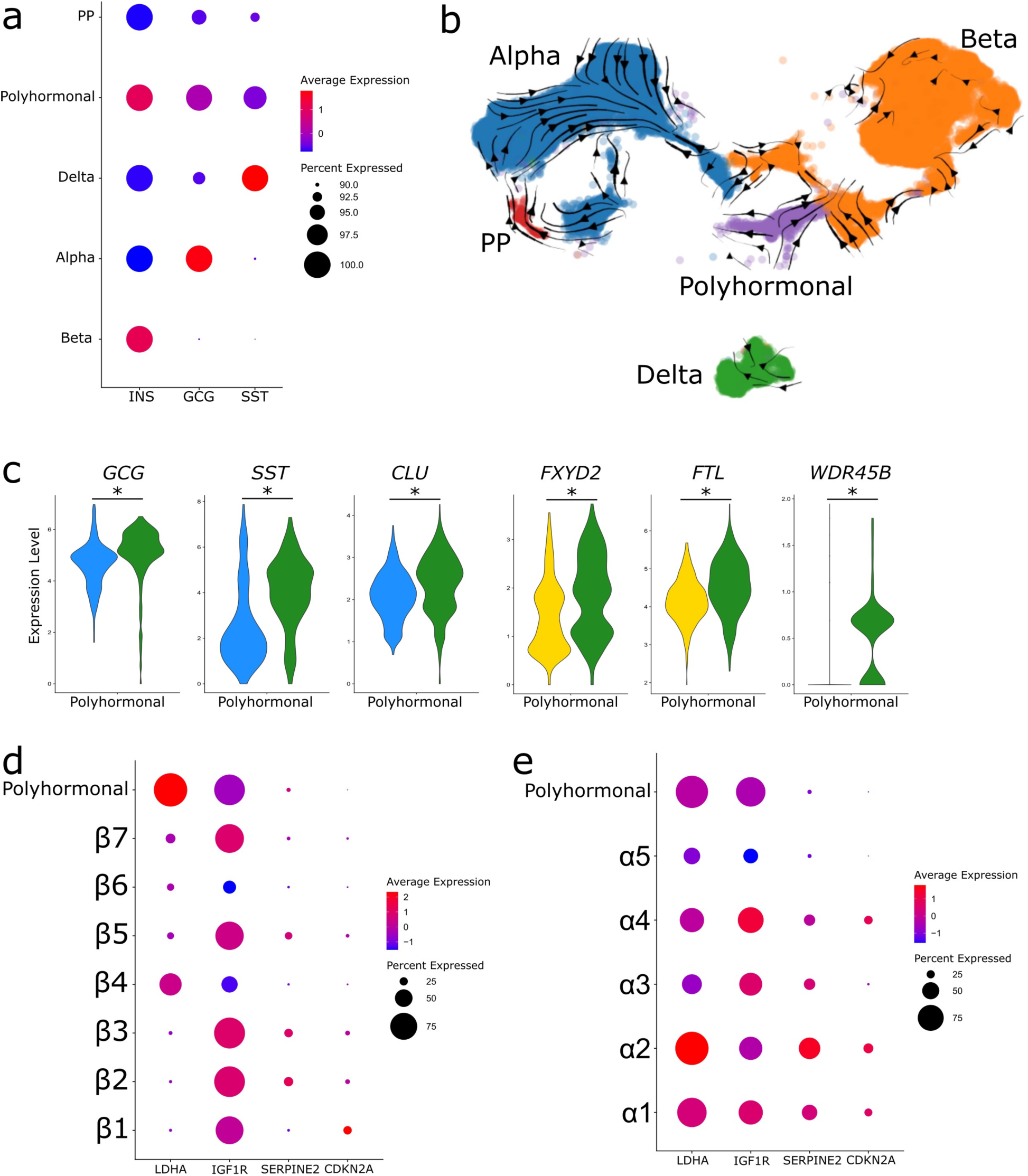
The polyhormonal cluster is transcriptionally distinct from α and β cells. **a**) Dot plot showing average expression of *INS, SST* and *GCG* in endocrine cell types. **b**) RNA Velocity trajectories from scVelo projected onto endocrine cell types in UMAP space. **c**) Split-violin plots showing representative calcium-regulated and glucose-regulated genes in the polyhormonal cluster. \**p*adjusted<0.05. **d**, **e**) Dot plots showing average expression of known markers of senescent β cells in the polyhormonal cluster compared to β1–β5 (**d**) and α1–α5 (**e**).

Third, polyhormonal cells have their own unique set of calcium-regulated and glucose-regulated genes that are not detected in cross-conditional comparisons in other cell types (Supplementary Table 6). Most importantly, we detected islet cells that expressed both *INS* and *GCG* mRNA using RNAscope fluorescent *in situ* hybridization (FISH) in sections of both *ex vivo* isolated human islets and human pancreas biopsies from multiple male donors, including the donors whose islets were used for the scRNA-seq libraries (Supplementary Fig. 8). Overall, we are confident that polyhormonal cells are a real population within the adult human islet, distinct from α or β cells that make up the majority of endocrine cells.

To investigate whether polyhormonal cells could be an immature endocrine cell population, we compared the average expression of known markers of mature α or β cells between polyhormonal cells, α cell clusters, and β cell clusters. Compared to α cell clusters α1-α5, polyhormonal cells express lower levels of α cell markers such as *GCG, TTR,* and *IRX2,* but similar levels of *ALDH1A1, CRYBA2, TM4SF4,* and *LOXL4* as α1 or α4 (Supplementary Fig. 7d). Compared to β cell clusters, polyhormonal cells expressed similar levels of *INS* as β2, β3, or β7, but lower levels of almost all other mature β cell markers such as *IAPP, MAFA,* and *ERO1B* (Supplementary Fig. 7e). In addition, we performed RNA velocity analysis on the endocrine cell types to investigate if the polyhormonal cluster or any other cluster could potentially be transitioning cell fates to a different cell type. Strikingly, the trajectories within the polyhormonal cluster were oriented towards both α and β cells, while there were minimal trajectories within the β and δ cells (Fig. 5b). These results show that compared to mature α or β cells, polyhormonal cells are immature at the transcriptional level, but it’s possible that some of these polyhormonal cells have the potential to become mature α or β cells.

Given the age of the sequenced donors, an additional possibility was that polyhormonal cells were senescent cells^28–30^. we compared the average expression of previously identified senescence markers *IGF1R, CDKN2A,* and *SERPINE2* across α, β and polyhormonal clusters. None of the three senescence markers showed an elevation in the polyhormonal cells compared to α or β clusters at the mRNA level (Fig. 5d,e). Within β cells, only less than 25% of cells in any cluster even expressed *CDKN2A* and *SERPINE2,* but most clusters except β4 and β6 expressed *IGF1R* (Fig. 5d). Within α cells, less than 50% of α1-α4 clusters expressed *CDKN2A* and *SERPINE2,* while polyhormonal cells did not (Fig. 5e). In addition to senescence markers, we found that the β cell “disallowed” gene *LDHA*^31^ was robustly expressed in the polyhormonal cells when compared with β cells, but not α cells (Fig. 5d,e).

As mentioned above, we determined the calcium- and glucose-regulated genes for the polyhormonal cluster and compared with those in α and β cells (Fig. 5c, Supplementary Table 6). We found 485 calcium-regulated and 1,001 glucose-regulated genes in the polyhormonal cluster, which is several times the number of regulated genes in any other cell type (Supplementary Table 6). The majority of these genes were unique to polyhormonal cells when compared with α or β cells (Supplementary Fig. 9a,b). Only 44 out of 453 calcium-induced genes and 29 out of 32 calcium-suppressed genes were also regulated in α and β cells. 39 out of 933 of the glucose-induced genes were also regulated in β cells, and 61 out of 68 genes that had higher expression in the Low condition (vs Positive) were unique to polyhormonal cells.

From these results, we can conclude that polyhormonal cells are a distinct population in the adult human islet, but the biological role of these cells is yet to be confirmed. They may be less mature than α or β cells at the transcriptional level, but express a large number of unique calcium- and glucose-regulated genes, as well as the β cell “disallowed” gene *LDHA*.

### PCDH7 is a marker of a novel β cell subtype with elevated function

Given the clearly different magnitudes of response to calcium signalling between β cell clusters, we reasoned that a method of identifying some of these clusters with higher calcium response could be useful in the future to study different β cell subtypes. Since clusters β1-β3 expressed the most calcium-regulated genes and activity-regulated genes, we decided to look for cell surface markers that could be used to enrich for this more calcium-responsive population. We found that protocadherin 7 (PCDH7), a transmembrane protein and member of the cadherin superfamily, was expressed in mostly β cells (Fig. 6a). Within the β cell clusters, *PCDH7* expression was particularly enriched in the β1-β3 and β7 clusters, while it was absent in the potentially immature β4 and β6 clusters (Fig. 6b). In a previous study, *PCDH7* was noted as one of the many upregulated DEG in *CD9*-negative β cells with elevated GSIS, but was not explored further as an independent marker of this elevated function^32^. Therefore, we hypothesised that PCDH7 alone could be a marker of mature β cells with elevated function.

**Figure 6.**
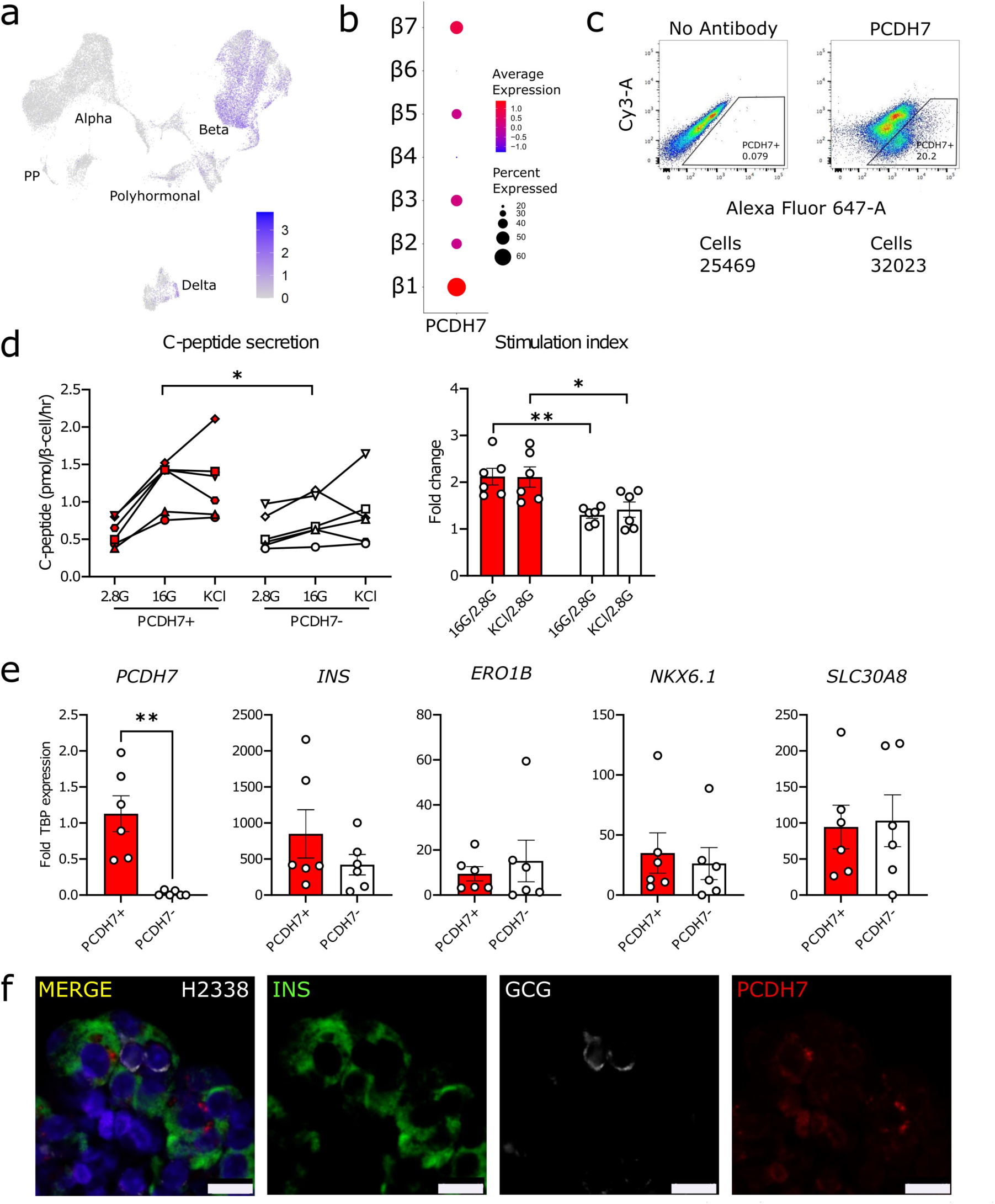
PCDH7 marks β cells with enhanced GSIS. **a**) UMAP plot of *PCDH7* expression in endocrine cells. **b**) Dot plot showing *PCDH7* expression in β cell clusters. **c**) Sorting results for isolating PCDH7^+^ cells from human islets. **d**) C-peptide concentrations from a static GSIS performed on reaggregated PCDH7^+^ and PCDH7^−^ β cells from donors R366, R367, R369, H2330, H2337, and H2338, shown as separate symbols. Cells were exposed to 2.8mmol/l glucose (2.8G), 16mmol/l glucose (16G), and 40mmol/l KCl (KCl). **e**) Key β cell maturity gene expression levels in PCDH7^+^ and PCDH7^−^ sorted cells. **f**) Immunostaining for INS (green), PCDH7 (red), GCG (white) and DAPI (blue) in a representative islet section of donor H2338. Scale bar, 10 μm. \**p*<0.05 and ** *p*<0.01.

Based on *PCDH7* expression levels, we divided all β cells in the dataset into *PCDH7*-high and *PCDH7*-low cells. Out of the 14,949 β cells, 4,886 (32.7%) cells were classified as *PCDH7*- high. A DEG analysis comparing *PCDH7*-high cells vs *PCDH7*-low cells revealed 311 genes that were enriched in *PCDH7*-high cells (Supplementary Table 7). The second-most enriched gene in *PCDH7*-high cells was *SPP1,* which was identified as a potential marker for the β1 cluster (Fig. 2c). This supports the finding that *PCDH7* was highly expressed in the β1 cluster, one of the most calcium-responsive β cell clusters. To validate whether *PCDH7*-high cells were also functionally different from *PCDH7-*low cells, human islets were sorted using an anti- PCDH7 antibody. Approximately 20% of the sorted human islet cells expressed cell surface PCDH7, which was a similar proportion to the *PCDH7-*high cells found in the scRNA-seq dataset (Fig. 7c). The sorted PCDH7^+^ and PCDH7^−^ cells were re-aggregated into spheroids, and their function was assessed with secreted C-peptide measurements during a static GSIS. PCDH7^+^ cells showed roughly two-fold higher GSIS stimulation index compared with PCDH7^−^ cells (Fig. 7d). This functional difference was not the result of differences in maturity state, indicated by the similar levels of *INS, ERO1B, NKX6-1,* and *SLC30A8* between PCDH7^+^ and PCDH7^−^ cells (Fig. 7e). Finally, immunostaining for PCDH7 in human islet sections showed that PCDH7 protein was present on the cell membrane of INS-positive β cells, but was absent in GCG-positive α cells (Fig. 7f). These results show that the cell surface protein PCDH7 is a marker of β cells that have a higher transcriptional response to calcium signalling and an enhanced GSIS response.

## Discussion

Here we used a multi-condition human islet scRNA-seq dataset to identify calcium-regulated genes in adult α, β and δ cells. We also showed distinct clusters of polyhormonal cells that express their own unique calcium-regulated profile, and histologically validated these cells in human islets. Finally, we found that the cell surface protein PCDH7 is a novel marker of the most calcium-responsive β cells transcriptomically, and these cells also display enhanced function. Overall, our study demonstrates that correlating calcium-regulated gene expression to function in islet cells can be used to reveal functional heterogeneity using scRNA-seq data.

A limitation of this study is the low number of donors used to generate the single-cell dataset. We specifically used islets from healthy male donors and tried to match donor biometrics (e.g. age, BMI, sex) as much as possible to reduce biological variation within the data. However, this did limit the total number of islet preps included in the experiment, and even with the matched donors, there were poorly integrated clusters that had to be removed after subsetting and reclustering the endocrine cells. The removal of poorly integrated clusters ensured that the identification of calcium-regulated genes was a good representation of all biological replicates, and the identity of the genes were not disproportionately affected by a single donor.

One unexpected finding was the regulation of *INS* expression in non-β cell populations. *INS* was detected as a calcium-regulated gene in two out of five α cell clusters and in δ cells, albeit at lower levels overall compared with β cell clusters. One possibility is that the non-physiological stimulatory conditions could have led to abnormal expression and regulation of *INS* in α and δ cells. While α cells can respond to high glucose, their activation and glucagon secretion would normally be inhibited in a hyperglycaemic environment due to paracrine signalling from β cells and δ cells^33–36^. However, we exposed the islets to high glucose and directly depolarised all populations, which would not occur in physiological conditions but might occur in pathophysiological conditions like diabetes. It is possible that under these conditions, non-β cells can express a low level of *INS*. Whether this is translated to the protein level is unknown, but could have implications in the aetiology of type 2 diabetes.

While the goal was not to specifically study rare cell populations, we found cells that expressed *INS, GCG*, and *SST.* Previous studies have also found islet cells that express two or even three characteristic endocrine genes^10,12,37–39^. We did not observe any progenitor gene expression, so it is unlikely that this resulted from dedifferentiation of mature cells. In previous studies, there have been very few polyhormonal cells relative to the overall dataset. Within our dataset, we observed a distinct cluster of polyhormonal cells that clustered away from all other endocrine cell types. While the biological role of these cells within the islet is unknown, polyhormonal cells express large, unique sets of hundreds of calcium-regulated genes. It is possible that polyhormonal cells could be immature cells that have a hyperactive transcriptional response to calcium. In the future, it would be ideal to identify a marker specific for this population to isolate the cells directly from human islets for closer study. This could be difficult without a large supply of human islets, because polyhormonal cells make up less than 4% of all endocrine cells, according to the scRNA-seq.

When we did attempt to find any rare β cells that were previously established, such as virgin β cells, hub β cells and senescent β cells^23,28,40,41^, there was no single cluster that perfectly aligned with published gene expression profiles of these rare populations. While we found the clusters β4 and β6 had lower expression of many key β cell genes, these clusters had the highest expression levels of *INS* (Supplementary Fig. 7e). Even in α cells, α5 had reduced expression of α cell marker genes and the lowest number of calcium-regulated genes, but had the highest levels of *GCG* expression (Supplementary Fig. 7d). This could indicate a trade-off between expression of hormone-encoding genes and calcium responsiveness.

In summary, our study demonstrates that calcium-regulated transcriptional profiles highlight differences in islet cell function and maturity. While mRNA expression only provides one view of the complex biology within the islet, we hope our dataset demonstrates the value in using transcriptomic approaches to study islet heterogeneity.

## Acknowledgements

We thank members of the Lynn laboratory and the Canadian Islet Research and Training Network for their helpful discussions and technical support. We thank Tatsuya Kin, James Lyon and Joss Manning Fox for provision of human islets. We thank the families of the human islet donors who made this research possible.

## Data availability

All data needed to evaluate the conclusions in the paper are present in the paper and/or the Supplementary. Requests for resources and further information should be directed to the lead contact, Francis C. Lynn. Sequencing data is available at NCBI Gene Expression Omnibus (GSE196715). A searchable and user-friendly format of the data in this study is available at https://lynnlab.shinyapps.io/Hislet_2023/.

## Contribution statement

JSY and FCL designed the overall research. JSY, MYYL, SS and CN performed experiments. JSY, SS, MYYL, JV, KZ, and HW analysed data. JSY, SS, JV, HW and FCL wrote the manuscript. JV designed and made the website. All authors read and edited the manuscript before giving final approval of the version to be published. . FCL is responsible for the integrity of the work as a whole.

## Abbreviations

α: Alpha
β: Beta
δ: Delta
CAMK: Calmodulin-dependent protein kinase
CaN: Calcineurin
CREB: cAMP response element-binding protein
DEG: Differentially expressed genes
GCG: Glucagon
GSIS: Glucose-stimulated insulin secretion
IEG: Immediate early genes
INS: Insulin
KRBH: Krebs-Ringer Bicarbonate HEPES
NFAT: Nuclear factor of activated T cells
OxPhos: Oxidative phosphorylation
PCDH7: Protocadherin 7
PP: Pancreatic polypeptide
qPCR: Quantitative PCR
scRNA-seq: Single-cell RNA sequencing
SST: Somatostatin
TCA: Tricarboxylic acid
UMAP: Uniform Manifold Approximation and Projection

## Supplementary Materials

**Includes:**

Supplementary Methods: antibodies and primers

Supplementary Figures S1-S9

Supplementary Tables S1-S7 (separate excel file)

## Supplementary Methods: Antibodies and Primers

### Antibodies used for immunofluorescence staining

| Primary or Secondary | Antibody | Dilution | Manufacturer |
| --- | --- | --- | --- |
| Primary | guinea pig anti-insulin | 1:500 | Dakocytomation, Denmark (A0564) |
| Primary | rabbit monoclonal anti-glucagon | 1:100 | Abcam, United Kingdom (ab92517) |
| Primary | mouse monoclonal anti-glucagon | 1:2000 | Sigma-Aldrich, USA (G2654-100UL) |
| Primary | rabbit polyclonal anti-IGFBP2 | 1:200 | Thermo Fisher Scientific, USA (15699-1-AP) |
| Primary | rabbit monoclonal anti-Neuropeptide Y | 1:400 | Cell Signaling Technology, USA (11976S) |
| Primary | mouse monoclonal anti-PCDH7 | 1:100 | Abcam, United Kingdom (ab139274) |
| Secondary | donkey anti-guinea pig FITC | 1:450 | Jackson ImmunoResearch, USA (706-096-148) |
| Secondary | donkey anti-rabbit FITC | 1:450 | Jackson ImmunoResearch, USA (706-166-148) |
| Secondary | donkey anti-mouse Cy3 | 1:250 | Jackson ImmunoResearch, USA (715-166-150) |
| Secondary | donkey anti-guinea pig Cy3 | 1:250 | Jackson ImmunoResearch, USA (711-096-152) |
| Secondary | goat anti-rabbit Alexa Fluor 647 | 1:200 | Invitrogen, USA (A27040) |

### Primers used for qPCR

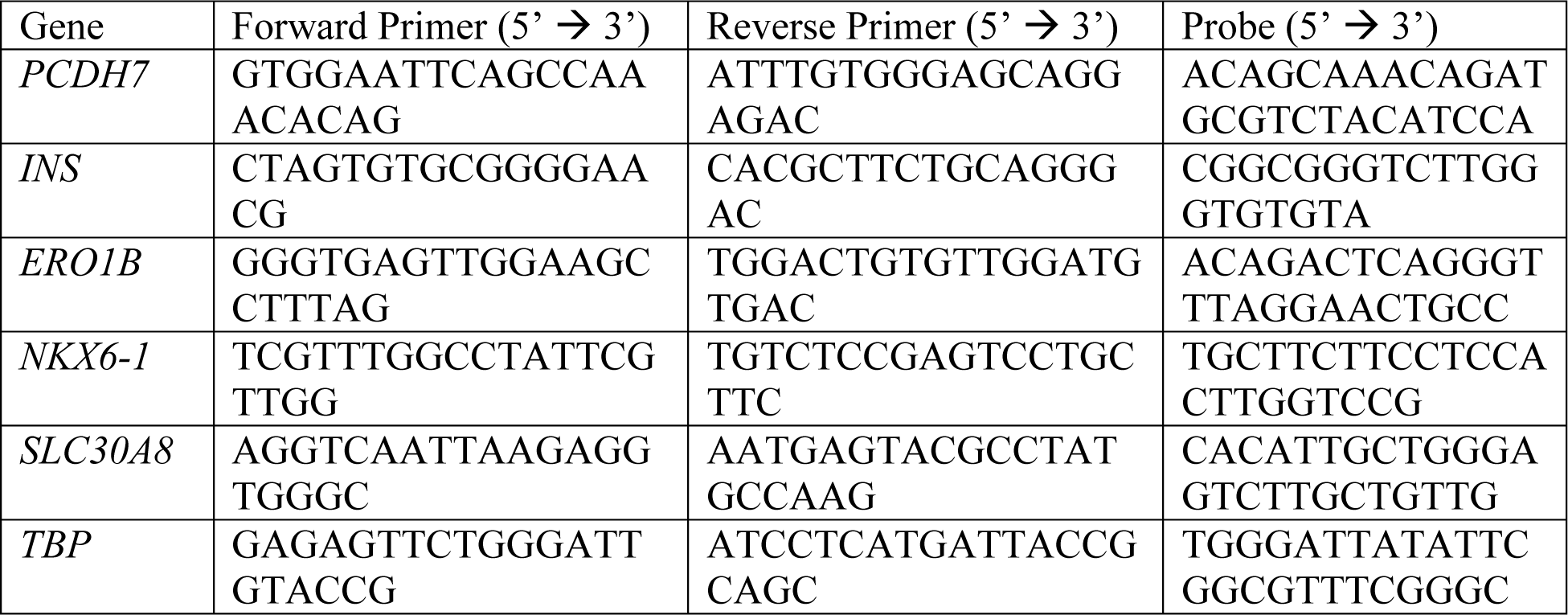

