## Supplementary Figures for "Calcium-dependent transcriptional profiles of human pancreatic islet cells reveal functional diversity in islet subpopulations"

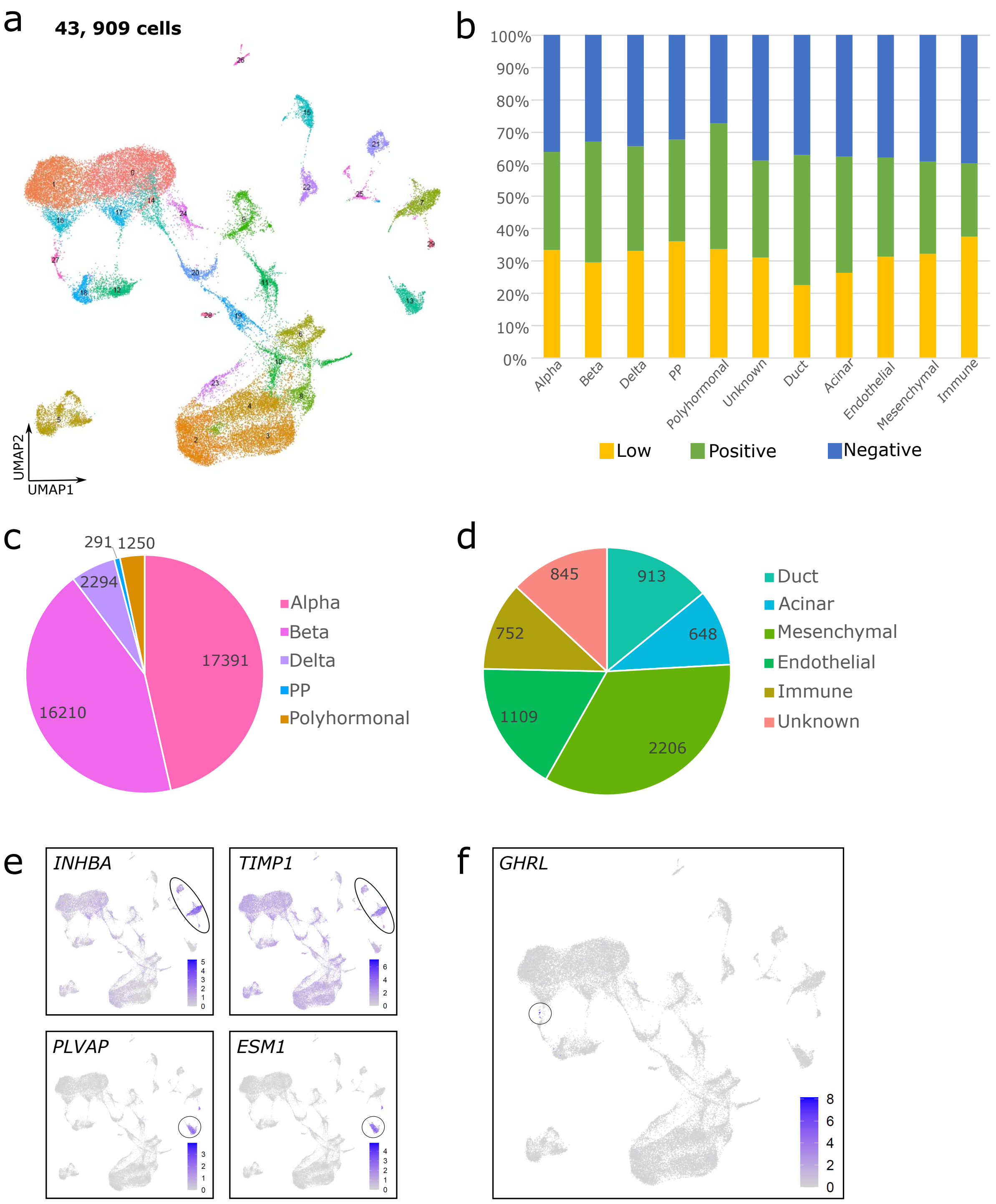


**Supplementary Figure 1. Human islet dataset pre-removal of cells.** a) UMAP plot of 43,909 human islet cells as 30 clusters. b) Proportions of cells from Low, Positive, and Negative conditions in each cell type. c) Cell numbers in each endocrine and d) non-endocrine cell type out of 43,909 cells. e) UMAP plots of mesenchymal marker genes *INHBA* and *TIMP1*, and endothelial marker genes *PLVAP* and *ESM1*. f) UMAP projection of *GHRL.*

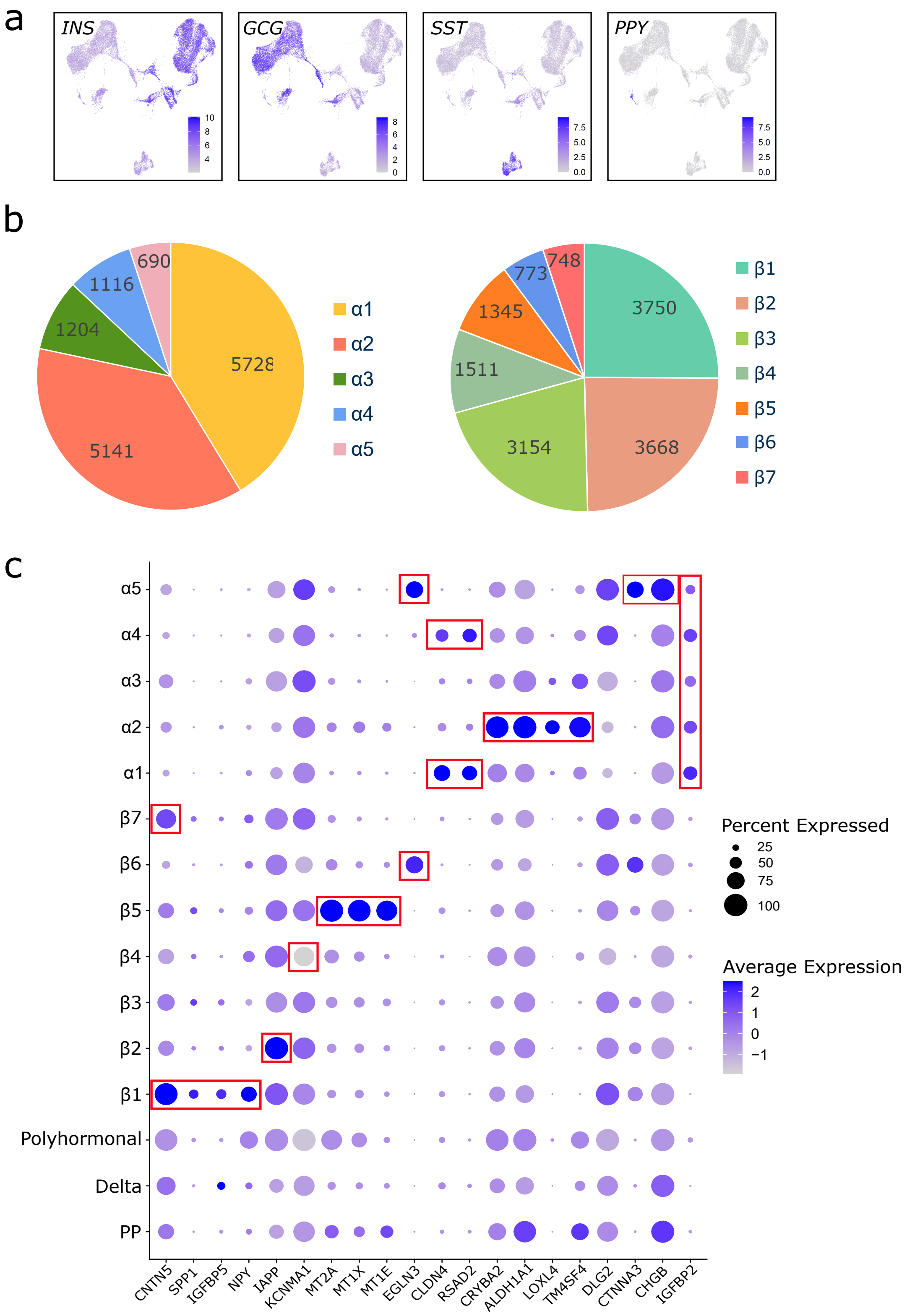


**Supplementary Figure 2. α and β cell clusters in the endocrine dataset.** a) UMAP plots of INS, GCG, SST, and PPY expression used to identify cell types. b) Breakdown of cell numbers in each α and β cell clusters. c) Dot plot showing the average expression for candidate cluster marker genes for each cluster. Dot size shows the proportion of cells in each cluster that expresses the genes on the X-axis. Dots highlighted in red boxes are leading candidate genes for each cluster.

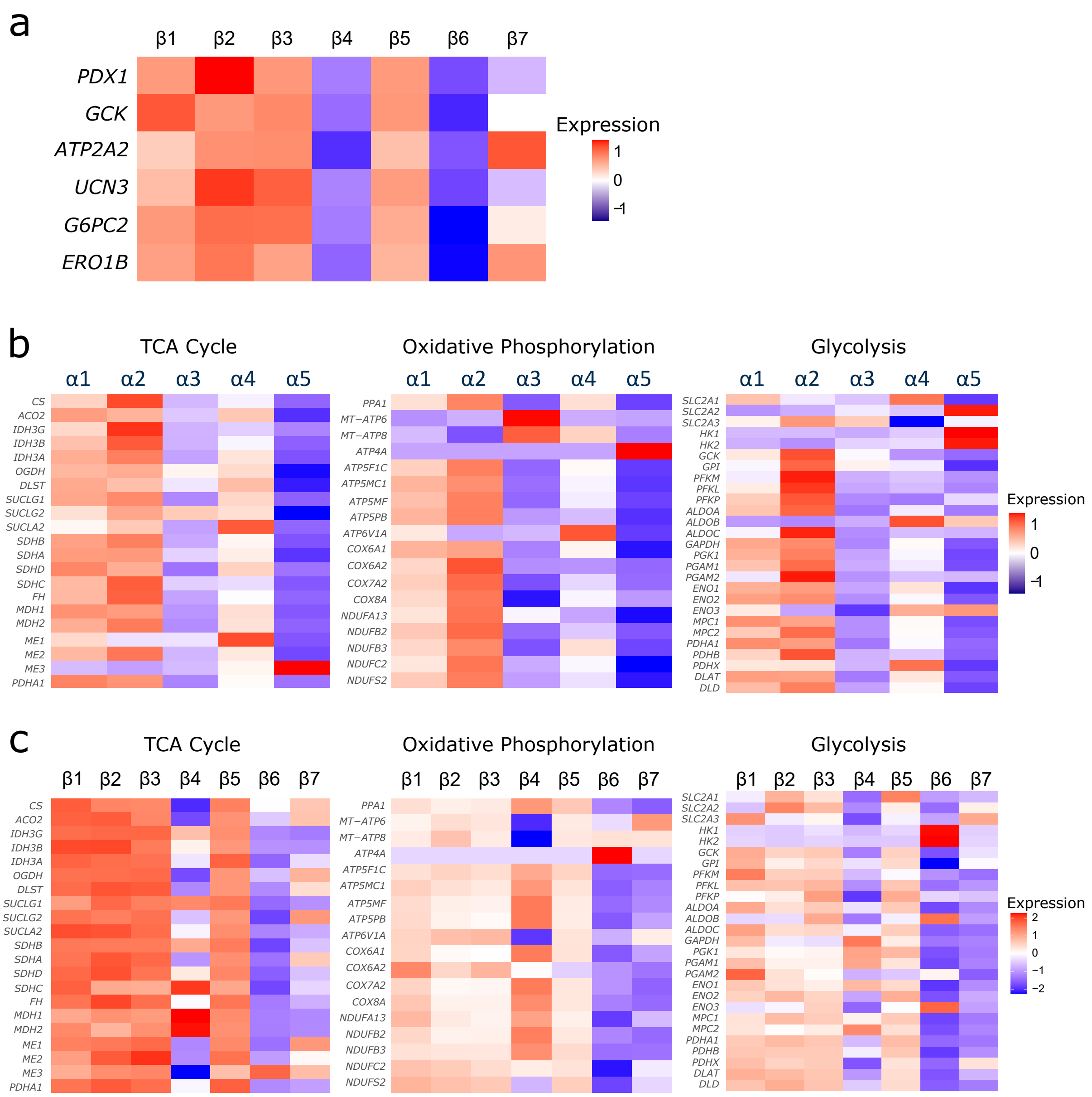


**Supplementary Figure 3. Maturity and metabolism gene expression profiles of β cell clusters.** a) Heatmap of average expression per cluster of key β cell maturity genes across clusters β1- β7. b) Heatmap of average expression per cluster of genes involved in TCA cycle, oxidative phosphorylation, and glycolysis across α1-α5 and c) β1- β7. Expression shown as a range of red (high) to blue (low) comparing each cluster to all other clusters.

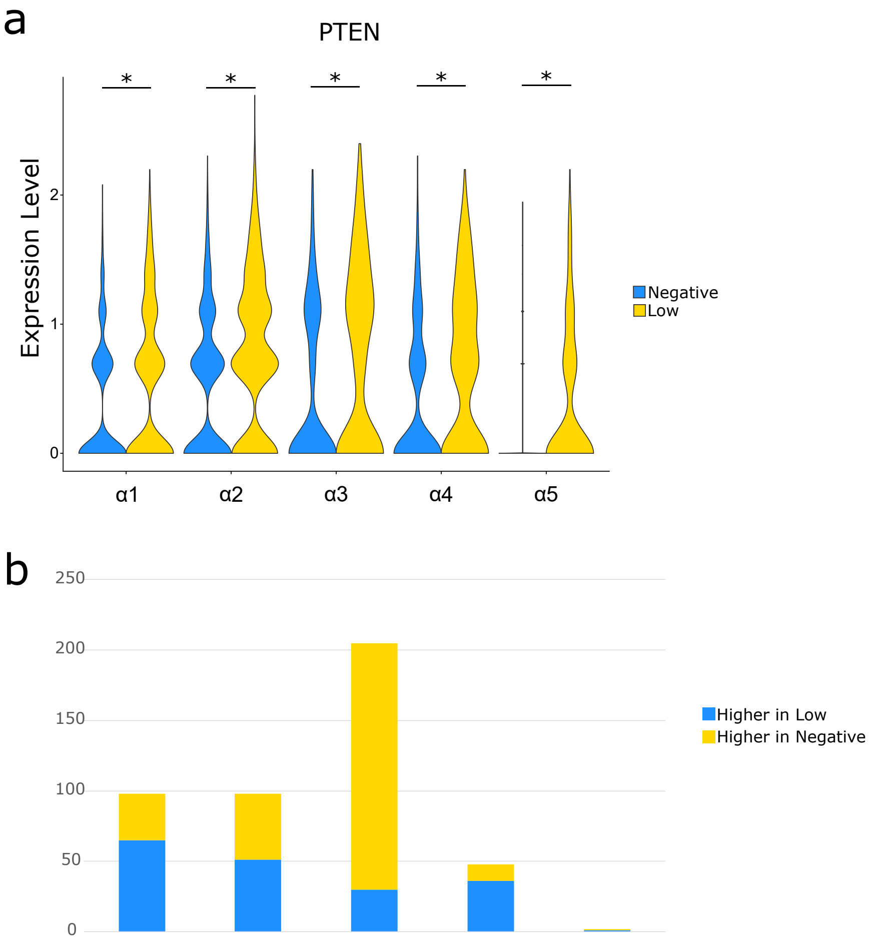


**Supplementary Figure 4. Comparing Low and Negative conditions in α cells.** a) Split violin plots showing expression of *PTEN* across clusters α1-α5 in the Low (yellow) and Negative (blue) conditions. *$p_{adjusted}<0.05$, Low vs Negative. b) Stacked bar graphs showing total numbers of genes expressed higher in the Low condition (yellow) vs higher in the Negative condition (blue) across clusters α1-α5.

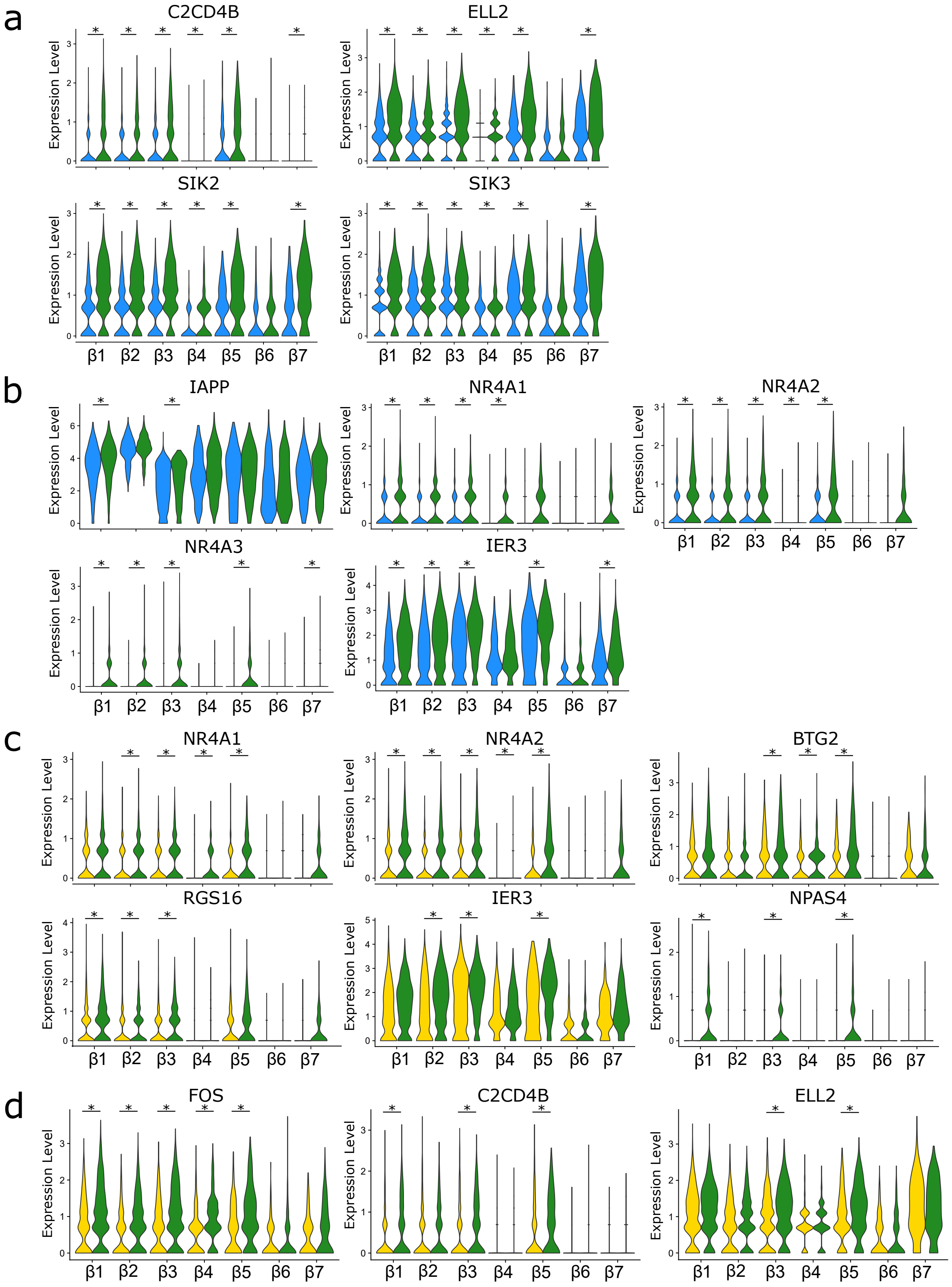


**Supplementary Figure 5. Calcium-regulated and glucose-regulated genes in β cells.** Split violin plots showing expression of a) calcium-regulated genes *C2CD4B, ELL2, SIK2, SIK3*, b) *IAPP, NR4A1, NR4A2, NR4A3*, and *IER3* across clusters β1- β7 in the Positive (green) and Negative (blue) conditions. *$p_{adjusted}$<0.05, Positive vs Negative. c) Split violin plots showing expression of glucose-regulated genes *NR4A1, NR4A2, BTG2, RGS16, IER3, NPAS4, d) FOS, C2CD4B,* and *ELL2* across clusters β1- β7 in the Low (yellow) and Positive (green) conditions. *$p_{adjusted}$<0.05, Positive vs Low.

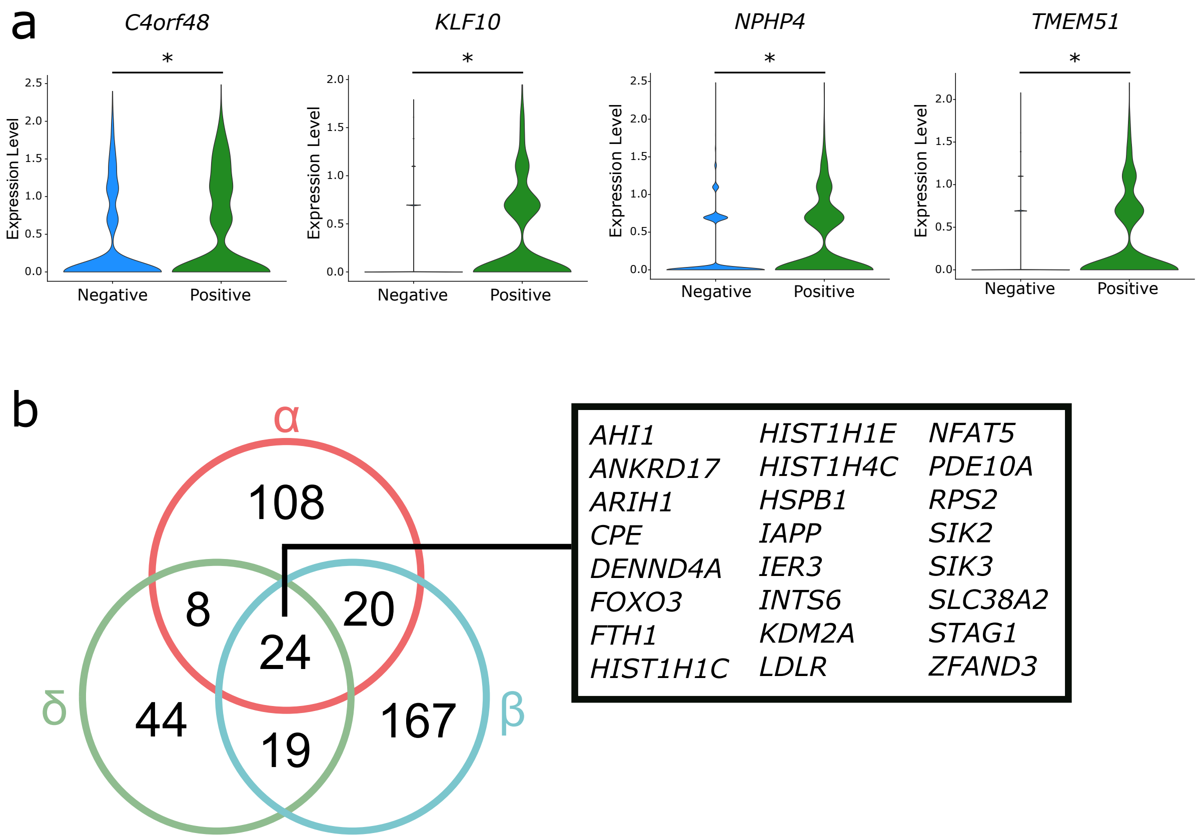


**Supplementary Figure 6. Calcium-regulated and glucose-regulated genes in α, β, and δ cells.** a) Split violin plots showing expression of genes *C4orf48, KLF10, NPHP4*, and *TMEM51* in δ cells in the Positive (green) and Negative (blue) conditions. $*p_{adjusted}<0.05$, Positive vs Negative. b) All glucose-regulated genes in α, β, and δ cells.

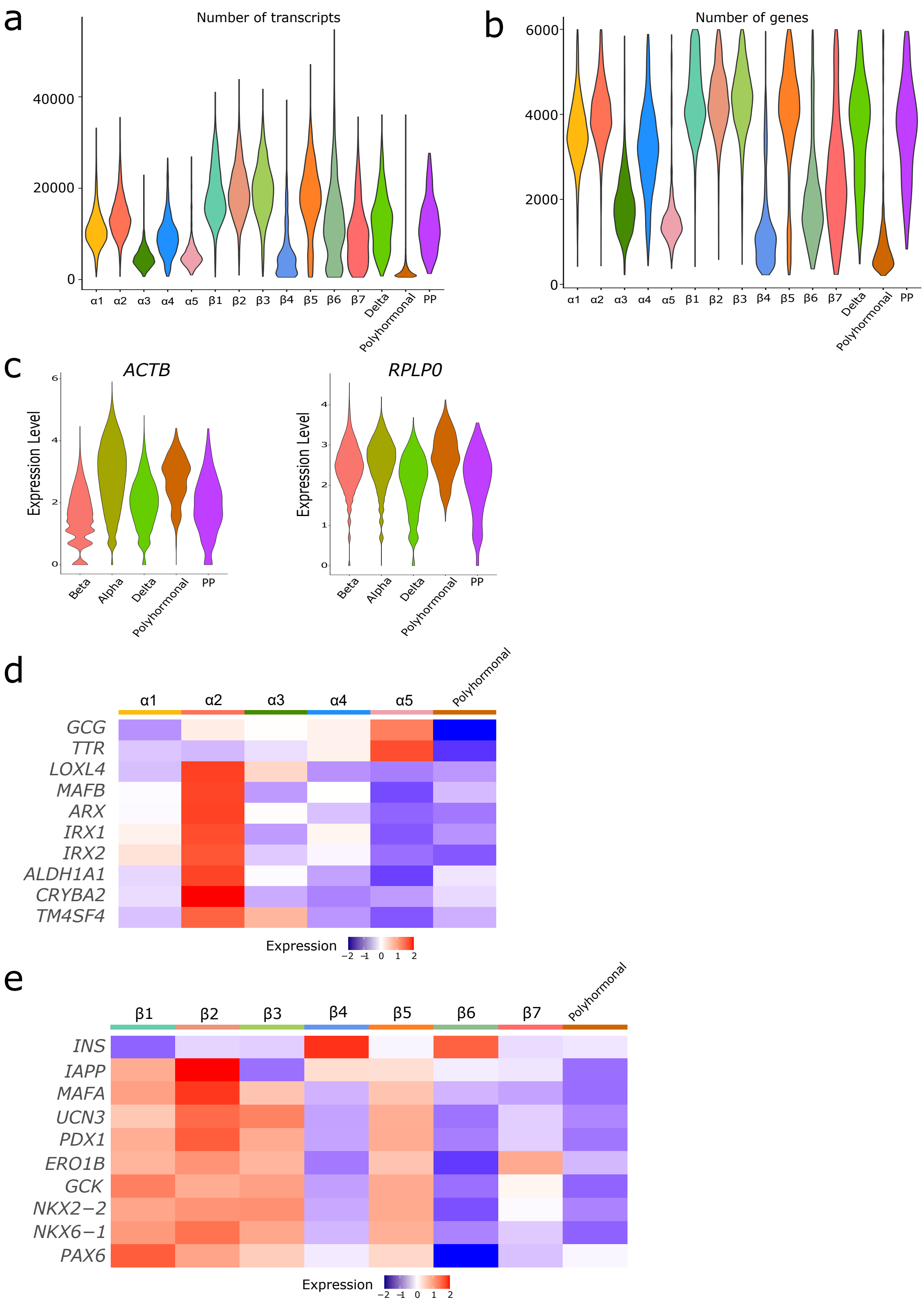


**Supplementary Figure 7. Comparing polyhormonal cells to α and β cells.** Violin plots of a) number of genes and number of transcripts in each cluster, and b) expression of housekeeping genes *ACTB* and *RPLP0* across all clusters. d) Heatmap showing average expression per cluster of α cell maturity genes and e) β cell maturity genes across the polyhormonal cluster, α cell clusters, and β cell clusters.

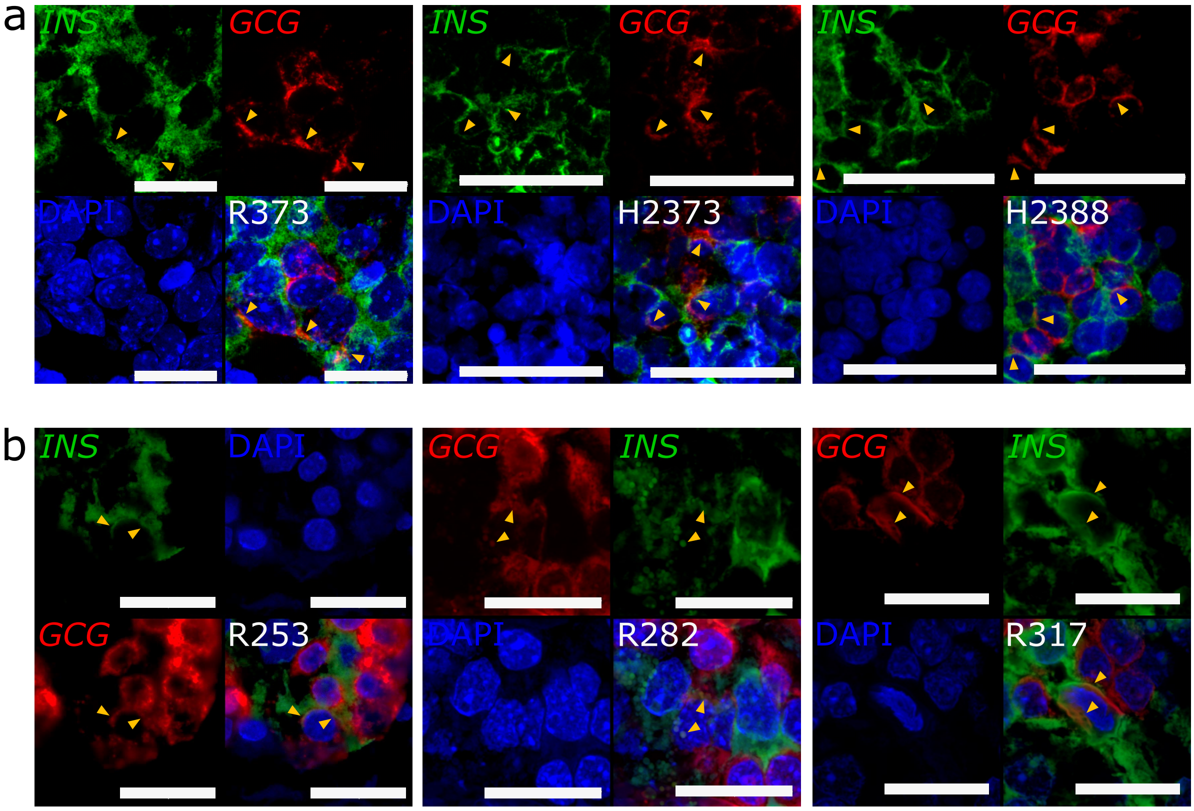


**Supplementary Figure 8. Histological validation of polyhormonal cells.** a) Representative images of *INS* (green) and *GCG* (red) mRNA detected by RNA FISH with DAPI nuclear stain (blue) in *ex vivo* human islet sections of donors R373, H2373, and H2388. b) Representative images of *INS* (green) and *GCG* (red) mRNA detected by RNA FISH with DAPI nuclear stain in islets within human pancreas biopsy sections of donors R253, R282, and R317. Arrowheads indicate regions of *INS* and *GCG* mRNA co-localization. Scale bars indicate 20um in R373, R253, R282, and R317 islets; 50um in H2373 and H2388 islets.

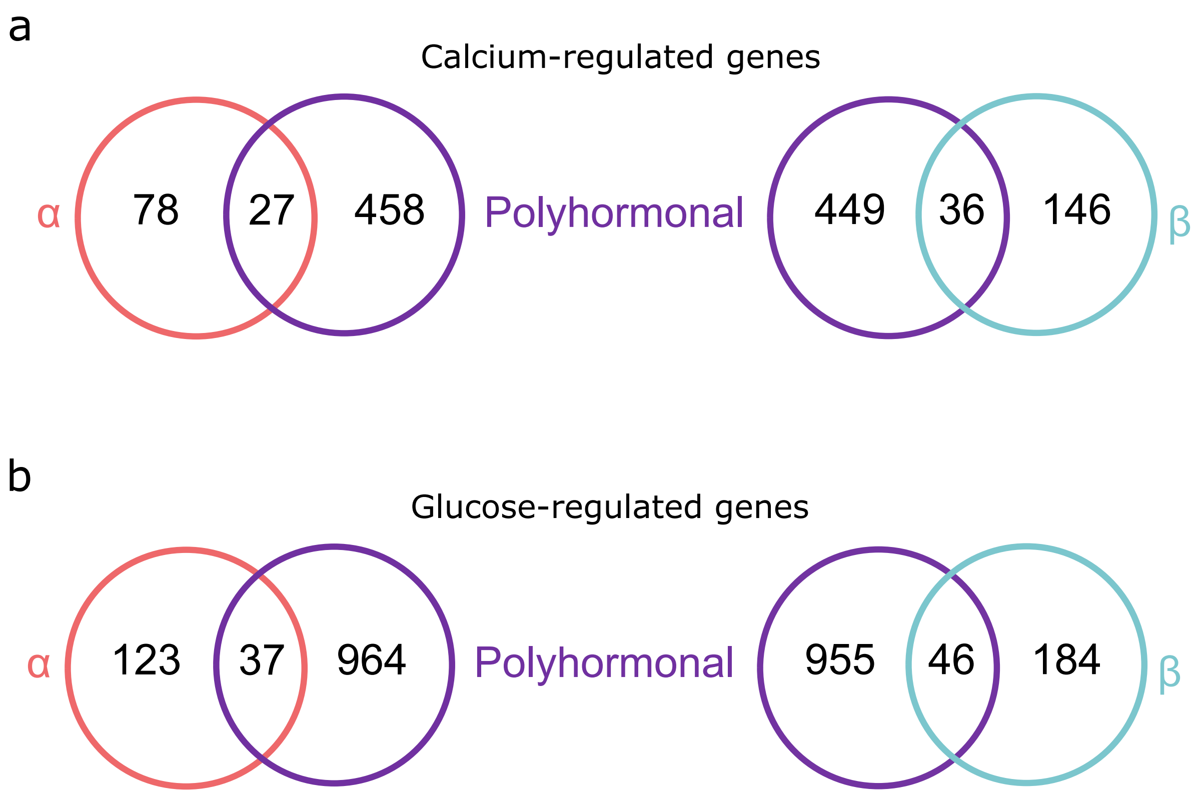


**Supplementary Figure 9. Comparing calcium-regulated and glucose-regulated genes in polyhormonal cells with α and β cells.** a) Numbers of calcium-regulated genes in polyhormonal cells vs. α and β cells. b) Numbers of glucose-regulated genes in polyhormonal cells vs. α and β cells.
